# Different stabilizing mechanisms but a common task-level aim in standing and walking

**DOI:** 10.1101/2025.11.24.690121

**Authors:** Yang Geng, Jaap H. van Dieën, Sjoerd M. Bruijn

## Abstract

Intrinsic mechanisms and feedback control act together to stabilize the body in upright tasks. In unperturbed standing and walking, their combined effects can be captured by lumped stabilization models, relating delayed center of mass position and velocity information to current ground reaction forces. If and how lumped stabilization parameters change across these tasks is unclear. We applied a stabilization model to estimate and compare stabilization between unperturbed standing and walking.

Fifteen healthy young participants (21± 4 yrs, 63 ± 9 kg, 1.70 ± 0.10 m, 13 females, 2 males) walked at 1.25 m/s for 5 minutes, and performed 3 different standing tasks: normal standing, unipedal standing, and step posture for 1 minute, repeated 5 times. Whole-body kinematics and ground reaction forces were collected and used to fit the stabilization model and estimate the effective delay and lumped gains. Only small differences were found between standing tasks. Model fits were significantly higher in standing than in walking. ML effective delays were significantly longer in walking than in standing, whereas AP delays were comparable. The stabilization gains varied significantly across tasks and directions. The lumped position gains in most tasks exceeded critical stiffness, except for the mean values in walking, which were lower than the critical stiffness. Lumped velocity gains were all at under-damped level. The ratio of lumped position to velocity gains in standing were consistently close to human body’s eigenfrequency 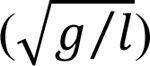 as predicted by the extrapolated center of mass concept. In walking, the ratio was close to the eigenfrequency during phases in which stabilization was significant.

Our findings suggest that stabilization is organized at the task level, with lumped effects of all stabilizing mechanisms acting to preserve a consistent weighting between position and velocity contributions across tasks, effectively regulating the CoM motion to follow a pendulum-like trajectory.

## 1. Introduction

Standing and walking are two fundamental motor behaviors in human daily life. Both require sophisticated neuromuscular control to maintain an upright body orientation against the effects of unpredictable perturbations which are amplified by gravity. Such stabilization is achieved through the coordination of intrinsic mechanisms and feedback control [1–3]. Intrinsic mechanisms provide immediate resistance to perturbations based on segmental inertia and passive tissue proprieties (i.e., stiffness and damping). In contrast, feedback control is adaptive and generates corrective responses selectively based on delayed sensory information [3–7]. While identifying these two stabilizing mechanisms separately is methodologically challenging [8], their combined effects determine the performance and robustness of these motor tasks [1, 3]. Insight into the contribution of both mechanisms requires separating the intrinsic mechanisms and feedback control, but characterizing their lumped effects offers an overarching and functional view of how stability is controlled [9].

Given the different temporal and biomechanical characteristics of standing and walking, we expect stabilizing mechanisms to be different in these tasks. Standing is a quasi-static behavior, during which stability can rely on constant intrinsic proprieties and time-invariant feedback control to maintain a ‘fixed’ body orientation. Walking is a dynamic, cyclic motion, during which the body experiences a phase-dependent instability. While the intrinsic mechanisms might provide some instantaneous and basic stability during the stance phase, feedback control must be dynamically modulated to deal with the time-varying stability requirements throughout the gait cycle [9, 10]. These fundamental differences suggest that the relative contributions of intrinsic and feedback mechanisms and the resulting lumped stabilizing effects may differ between standing and walking.

Compared to the relatively time-invariant intrinsic mechanisms, feedback control is context-dependent and adaptive. Many studies have addressed the neural mechanisms [5, 6, 9–14]. These studies commonly assume that sensory estimates of the task-level variable kinematics, i.e., center of mass (CoM), are used to modulate corrective responses (e.g., EMG, ankle torque, foot placement) to achieve stabilization [5, 10, 11, 13]. While these models provide valuable insight into the neural mechanisms in stability control, they have some limitations. First, these models are generally derived under specific tasks, i.e., either in standing [5, 11, 13] or walking [10, 14, 15], making it difficult to directly compare different motor tasks. Second, identification of feedback parameters usually requires perturbations and the measurements of physiological signals (e.g., EMG), which limits their application in studying natural, unperturbed behavior. Furthermore, focusing solely on feedback control fails to provide a comprehensive understanding of how overall stability is maintained and changes across tasks, specifically for those with fundamentally different dynamics.

Since all corrective actions will ultimately be reflected by the ground reaction forces [16, 17], the lumped stabilizing mechanisms can be represented by the relationship between current ground reaction forces and delayed CoM deviations. Based on this, van Dieen et al. [18] proposed a model to assess stabilizing control in unperturbed walking by fitting a linear relationship between the delayed center of mass kinematics and horizontal ground reaction forces. More recently, we refined this model by using the CoM position (PCoM) and velocity (VCoM) information as separate predictors instead of combined into a single predictor [19], which enabled us to estimate the position- and velocity-related control. We now have a framework to evaluate and compare lumped stabilization between standing and walking without external perturbations or physiological measurements. It should be noted that we have previously referred to these models as ‘feedback models’, however, because they also capture intrinsic mechanisms, we avoid this term here to avoid confusion and instead use the term ‘stabilization models’. These models can provide a better understanding of how the control system regulates overall stabilization based on preceding CoM kinematics, and how the parameters characterizing stabilization change across tasks.

The extrapolated center of mass (XCoM) concept, which equals the PCoM plus VCoM divided by the eigenfrequency (√*g*⁄*l*) of the body modeled as an inverted pendulum [20], couples the CoM position and velocity into a single metric.

According to this concept, stability is achieved by keeping the XCoM within the base of support (BoS) [20, 21]. This suggests that stabilization relies on coordinated control of CoM position and velocity, with the ratio of the control gains equal to the body’s eigenfrequency. While the lumped position and velocity gains themselves are expected to differ to meet specific stability requirements, their relative contribution, expressed as the ratio of position to velocity gains, may remain consistent across tasks if a common pendulum-like stabilization aim exists. Especially for walking, a phase-dependent task, whether and during which phases stabilization around a pendulum-like trajectory occurs remains an open question.

To address the above questions, we applied a model to quantify stabilization (‘stabilization model’) to unperturbed walking and typical standing postures (normal standing, unipedal, and step postures). Unipedal standing and step posture were selected as they are commonly observed during walking, while they provide a different base of support (BoS) and different structural stiffness compared with normal standing. Comparing these postures allows us to determine whether a consistent position-to-velocity control exists across different biomechanical contexts. Based on our previous findings [18, 19], we hypothesized that: 1) effective delays are longer in walking than in standing, because stabilizing effects become effective mainly after ground contact of the leading leg; 2) lumped gains are larger in walking than in standing postures, to compensate for larger delays (c.f. Foulger, Liu et al. [22]); 3) the ratio of lumped position to velocity gains is constant and close to the eigen frequency of the body modeled as an inverted pendulum.

## 2. Methods

### 2.1 Experimental design & data collection

Fifteen healthy young participants (21± 4 yrs, 63 ± 9 kg, 1.70 ± 0.10 m, 13 females, 2 males) participated in the experiment. The experimental procedure was approved by the ethics committee of the Faculty of Behavioural and Movement Sciences, Vrije Universiteit Amsterdam (number: VCWE-2023-170). Data collection started on September 24, 2024, and ended on November 11, 2024. All participants received oral and written explanations of the procedures and signed written consent before the experiment.

In this study, participants performed one walking trial and three different standing tasks. For walking, participants walked on an instrumented dual-belt treadmill (Motek-Force-link, Amsterdam, Netherlands) with belt speed set at 1.25 m/s (4.5 km/h) until a steady gait was achieved, followed by an additional 5 min walking trial. All participants reported that this speed felt natural without requiring excessive effort or causing fatigue. For standing tasks, participants stood on the instrumented dual-belt treadmill and were asked to perform: 1) Normal standing: stand upright with feet together. 2) Unipedal standing: stand on the preferred leg with the other leg in a position in which the participants could maintain stability. We tested the preferred leg only, because previously no differences were observed between standing on the dominant and non-dominant leg [23]. 3) Step posture: participants took a natural step as if walking normally at least three times until they confirmed to have reached the most comfortable step length and width, then maintained that position with body weight evenly distributed between both feet. This even distribution was selected as our pilot study showed no consistent differences between different weight-distributions. Foot positions were marked for consistency across trials. All standing tasks were kept for 1 min and repeated 5 times in random order. For each standing trial, participants faced forward and kept their arms naturally at their sides, trying to keep as still as possible. Sufficient rest time was given between tasks to avoid fatigue. Trials with stability loss (e.g., ground contact with the non-stance leg in unipedal standing, excessive arm movement) were discarded and repeated.

### 2.2 Data processing

Data collection and processing followed procedures from our previous study [19].

In short, we collected whole-body 3D kinematic data using Optotrak cameras (Northern Digital Inc., Waterloo, Ontario, Canada) and ground reaction force using force plates embedded in the treadmill (Motek-Force-link, Amsterdam, Netherlands), and processed these data using MATLAB (2025b, The MathWorks, Natick, US). Both kinematic and force data were low-pass filtered using a second-order Butterworth filter with a cut-off frequency of 10 Hz, and the force data were down-sampled to 50 Hz to match the kinematics sampling frequency. The whole-body center of mass position (PCoM) was calculated using a 3D linked segment model [24], and the center of mass velocity (VCoM) was calculated as the time derivative of PCoM. These variables were further processed differently for standing and walking.

For standing tasks, the mean PCoM was defined as the reference upright position, accordingly, PCoM was calculated as the deviation from this reference. The stabilization effort, represented by the corrected ground reaction forces (*F_corr_*), was obtained by removing the gravitational acceleration component from the total ground reaction forces (*F*_g*r*_), assuming the whole body moved as a single inverted pendulum (IP), in both ML and AP directions. The removal process is illustrated in Fig 1, with further details in [19]. For model fitting, we analyzed experimental data from seconds 6 to 50 of each trial. This interval was selected to exclude initial postural adjustments at the start and increased body sway at the end of the task, particularly in unipedal standing.

**Fig 1.**
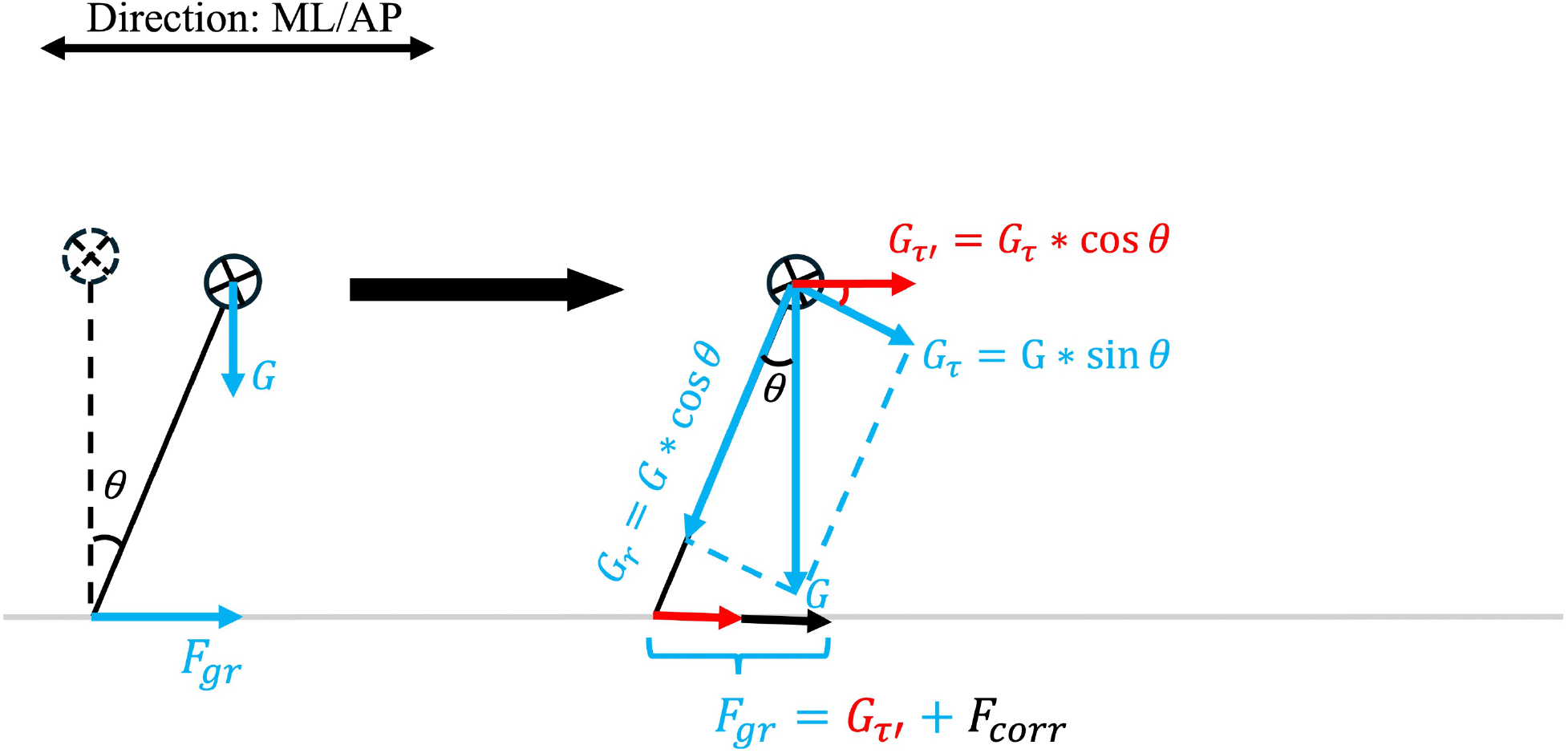
Removal of the gravity component. In this figure, the human body is modelled as a single inverted pendulum (IP). The left plot represents the IP states, with the dashed line indicating the reference upright state, while the solid line shows a representative IP state with a sway angle θ. In this representative state, the IP experiences the gravitational force (*G*) and generates a ground reaction force (*F_gr_*) in the ML/AP direction. Due to the inclination, the horizontal *F_gr_* includes a passive force resulting from gravity in addition to the ground reaction force generated by muscle activity (*F_corr_*). The right plot further explains how we calculated the corrected ground reaction force *F_corr_*. Gravity can be decomposed into two components: a centripetal force (*G*_r_), which equals *G ∗ cos θ*, and a tangential force (*G*_τ_), which equals *G ∗ sin θ*. The effect of *G*_τ_ in both horizontal directions (*G*_τ′_) can be calculated as *G_τ_ ∗ cos θ*. Finally, *F_corr_* is obtained by *F_corr_* = *F_gr_* − *G*_τ′_.

For walking, PCoM, VCoM and forces were time-normalized to the gait cycle (from right/left heel strike to subsequent right/left heel strike), yielding 100 samples per stride and the PCoM was re-referenced to the mean trailing foot position during the 15%∼45% phase of each stride. As the inverted pendulum (IP) model is too simple to fully represent the complex characteristics of walking [25], particularly during the double-stance phase, we tested whether removing the gravitational component for walking as used in standing could be applied. Thus, we fit the stabilization model using both total ground reaction forces (*F*_g*r*_) and corrected ground reaction forces (*F_corr_*) to decide whether the gravitational component could be reasonably removed. Results showed that using corrected ground reaction forces (*F_corr_*) led to a significantly decreased model fit in the ML direction (**Results 4.1**). Thus, for walking, we used total ground reaction forces to fit the stabilization model, which implies that any effects of gravity would contribute to residual error after fitting the model. The whole 300s walking data were used to fit the model.

### 2.3 Stabilization model

We fitted the stabilization model for standing and walking separately given their time-invariant and phase-dependent control.

For standing, equation (1) was used:

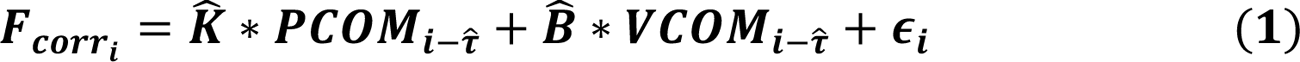

where K^ and B^ are the lumped position and velocity gains, *ε_i_* represents the residual error, *i* represents the current time point, τ^ represents the effective delay, which we allowed to range from 100 to 1000ms [11].

Based on our previous study [19], we evaluated the model’s goodness of fit using the coefficient of determination (*R*^2^), and estimated the lumped gains using ordinary least square regression, fitting *F_corri_* to the linear model specified in equation (1) for each delay (τ^). The effective delay was selected based on the best model fit (maximum *R*^2^), which is a common goodness of fit criterion in model fitting and regression analysis. The corresponding K^ and B^ were considered as the lumped position and velocity gains.

To test the reliability of these estimated parameters, we calculated the intra-class correlation coefficients (ICC) and SEM [26] for all standing tasks. Results supported using our approach to estimate the stabilization parameters for all standing tasks (**S1 Appendix**). The estimated effective delay (τ^), and lumped gains (K^and B^) of all trials were averaged over trials for statistical analysis.

For walking, equation (2) was applied to estimate the stabilizing parameters:

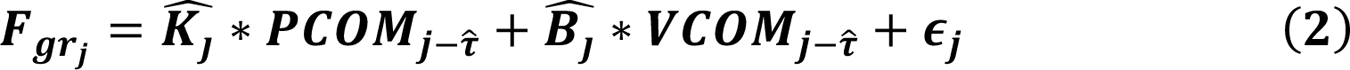

where (K^_j_ and B^_j_)represents the phase-dependent lumped position and velocity gains, *ε*_j_ represents the residual error, *j* represents the current phase (from 51% to 100% gait cycle), τ^ represents the effective delay ranging from 1% to 50% gait cycle. Model fit in walking was estimated using the same method as in standing. However, the effective delay could not be simply determined as the delay at which the best model fit was obtained. During walking, the center of mass signal exhibits relatively low variance during certain phases. As a result, the numerically optimal fit in these phases does not reflect stabilization (e.g., the effective delay in the ML direction selected based on the best fit was less than 20ms, which is physiologically implausible). Thus, we selected the time lag that yielded the largest negative weighted sum of position and velocity gains averaged over the stance phase as the effective delay. For this selected effective delay, the lumped position and velocity gains that produced the largest (most negative) weighted sum were defined as the peak position and velocity gains. Because peak values represent stabilization at a specific phase of the gait cycle, we additionally calculated the mean values of model fit and lumped gains across the whole stance phase to represent overall stabilization during walking. Thus, both peak and mean gains were considered for walking, whereas only mean values were used for model fit when comparing walking with standing. For clarity, in the subsequent figures and tables, ‘WM’ denotes the mean values, whereas ‘WP’ donates the peak values.

We further performed normalization of lumped position and velocity gains based on their critical values, i.e., critical stiffness and critical damping. The critical stiffness (*K_cr_*) refers to the minimal stiffness required to stabilize the system, which was calculated by equation (3). The critical damping refers to the damping value under which the system returns to the target position without overshoot and was calculated by equation (4):

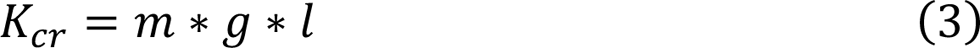

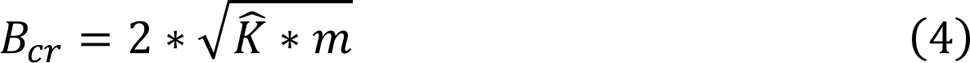

Where the *m* is body mass, *g* is the gravitational acceleration (9.81 *m*⁄*s*^2^), *l* is the center of mass height, K^ is the system’s current stiffness.

For the lumped position gain, the estimated linear form (*K^*) was first converted into its angular form by multiplying by *l*^2^, then normalized by dividing by the critical stiffness (*K_cr_*), yielding the normalized position gain (*K*^*_n_*). Similarly, for the lumped velocity gain, the estimated linear gain (*B^*) was first converted to rotational damping by multiplying by *l*^2^ and then normalized by dividing by the critical damping (*B_cr_*), yielding the normalized velocity gain (*B*^*_n_*). These normalized gains are unitless and describe how many times greater their rotational values are compared to their corresponding critical values.

To figure out whether the CNS modulates the lumped position and velocity gains based on a specific relationship as indicated by the concept of the extrapolated center of mass (i.e., based on the (angular) eigenfrequency), we calculated the ratio between the raw, unnormalized lumped position and velocity gains (K^/*B*B^), and further normalized this ratio relative to the eigenfrequency of the model represented as an inverted pendulum, by dividing by √*g*⁄*l*, represented as *R*^^^*_n_*. If indeed lumped position and velocity gains are modulated according to the extrapolated center of mass concept, we expect that *R*^^^*_n_* ≈ 1.

Simulations showed that, for standing, the identified stabilizing parameters are primarily determined by feedback control [19]. While intrinsic stabilization leads to systematic biases: under realistic combinations of intrinsic and feedback parameters, increased intrinsic stiffness and damping lead to a shorter delay and increased lumped position and velocity gains compared to the input feedback values. These biases should be taken into account when comparing the lumped values across tasks.

## 3. Statistical analysis

We examined the effects of Task and Direction on model fit, effective delay, and normalized gains by using a two-way repeated measures ANOVA with Task (walking, normal standing, unipedal standing, and step posture) and Direction (ML, AP) as within-participant factors to compare stabilizing parameters. Post-hoc pairwise comparisons with Bonferroni correction were conducted when a significant main effect of Task, Direction or an interaction was found.

For the ratio of position to velocity gains, as we were only interested in its deviation from the eigenfrequency, we used one-sample t-tests to compare the normalized ratio (*R*^^^*_n_*) against 1.

All statistical analyses were performed in MATLAB (2025b, The MathWorks, Natick, US). Statistical significance was set at α = 0.05. Results are presented as mean ± standard deviation (mean ± SD). Effect size for repeated measures ANOVA was evaluated using partial eta squared (*η*^2^*_p_*), with values of 0.01, 0.06, and 0.14 considered as small, medium, and large effects, respectively [27]. For the one-sample t-test, effect size was quantified using Cohen’s d, with absolute values of 0.2, 0.5, and 0.8 considered as small, medium, and large effects, respectively [28].

## 4. Results

Before the comparison between walking and standing, we first tested whether to use the total ground reaction forces or corrected ground reaction forces to fit the stabilization model to walking data.

### 4.1 Fitting model using total forces vs. corrected forces in walking

Model fit did not differ significantly between total ground reaction forces and corrected ground reaction forces in the AP direction (p=0.10), however, in the ML direction, using corrected ground reaction forces significantly decreased the model fit (p<0.001, Fig 2). In both directions, peak gains occurred in the early stance phase (∼60% gait cycle), regardless of using the total ground reaction forces or corrected forces.

**Fig 2.**
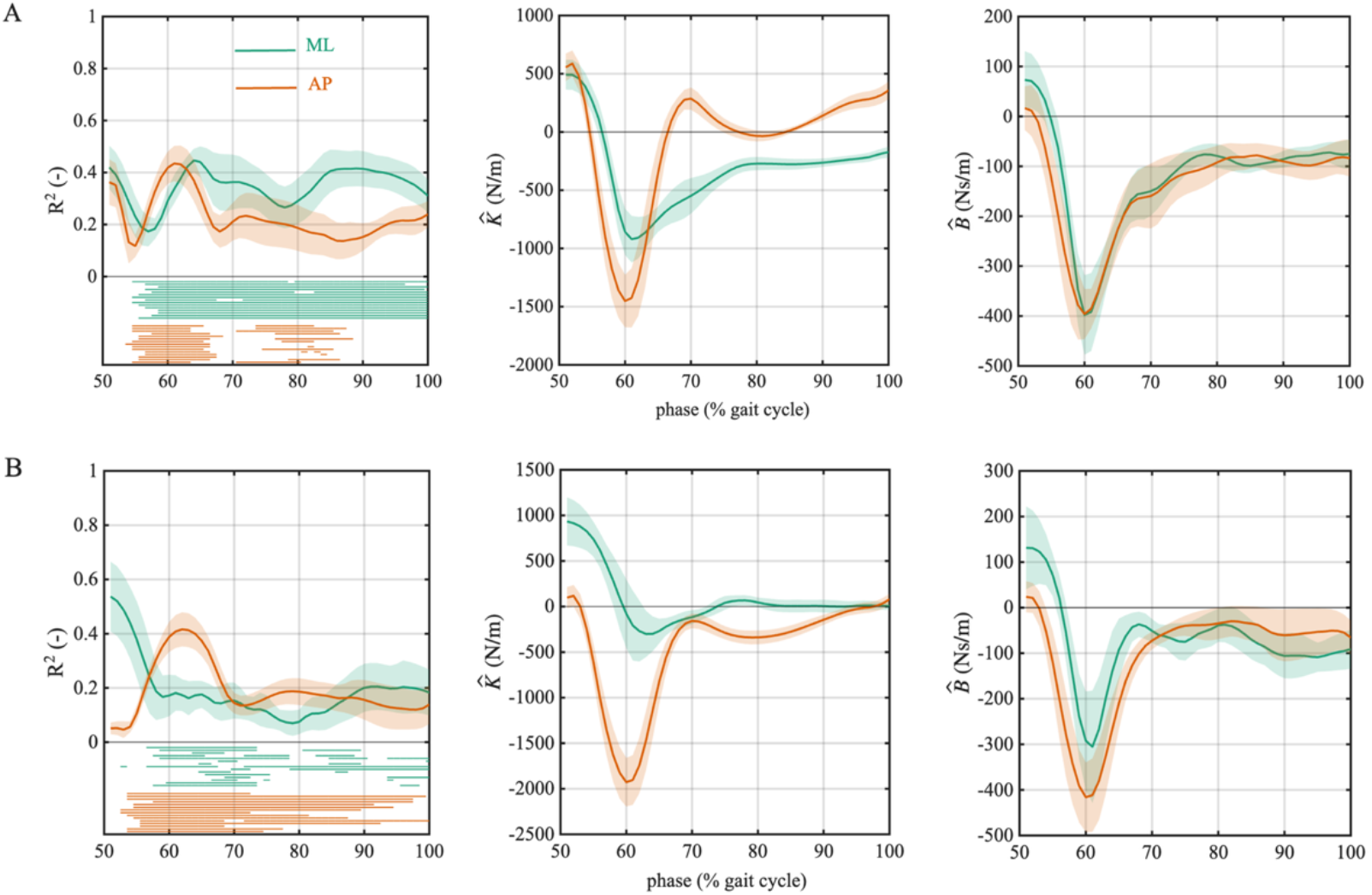
Stabilization models identified in walking with the total (panel A) and corrected ground reaction forces (panel B) as predicted variables. From left to right: model fit (*R*^2^), lumped position gains (*K^^^*) and velocity gains (*B^^^*) across the gait cycle. The green and orange lines represent the averaged results (over participants) in ML and AP direction, respectively. The shaded areas indicate the 95% confidence intervals. The horizontal lines in the left panels indicate phases in which the stabilization model was significant for each individual participant (p<0.05).

### 4.2 Comparison between tasks

The effective delay (τ^) and velocity gains (*B*^^^*_n_*) exhibited significant Task ∗ Direction interactions and significant main Task and Direction effects, whereas the model fit (*R*^2^) and position gains (*K*^^^*_n_*) exhibited interaction and Task effects but no significant Direction effects (Table 1, Fig 3). All observed effects had large effect sizes (*η*^2^*_p_*>0.14). These results showed that the stabilizing parameters differed across tasks and directions.

**Fig 3.**
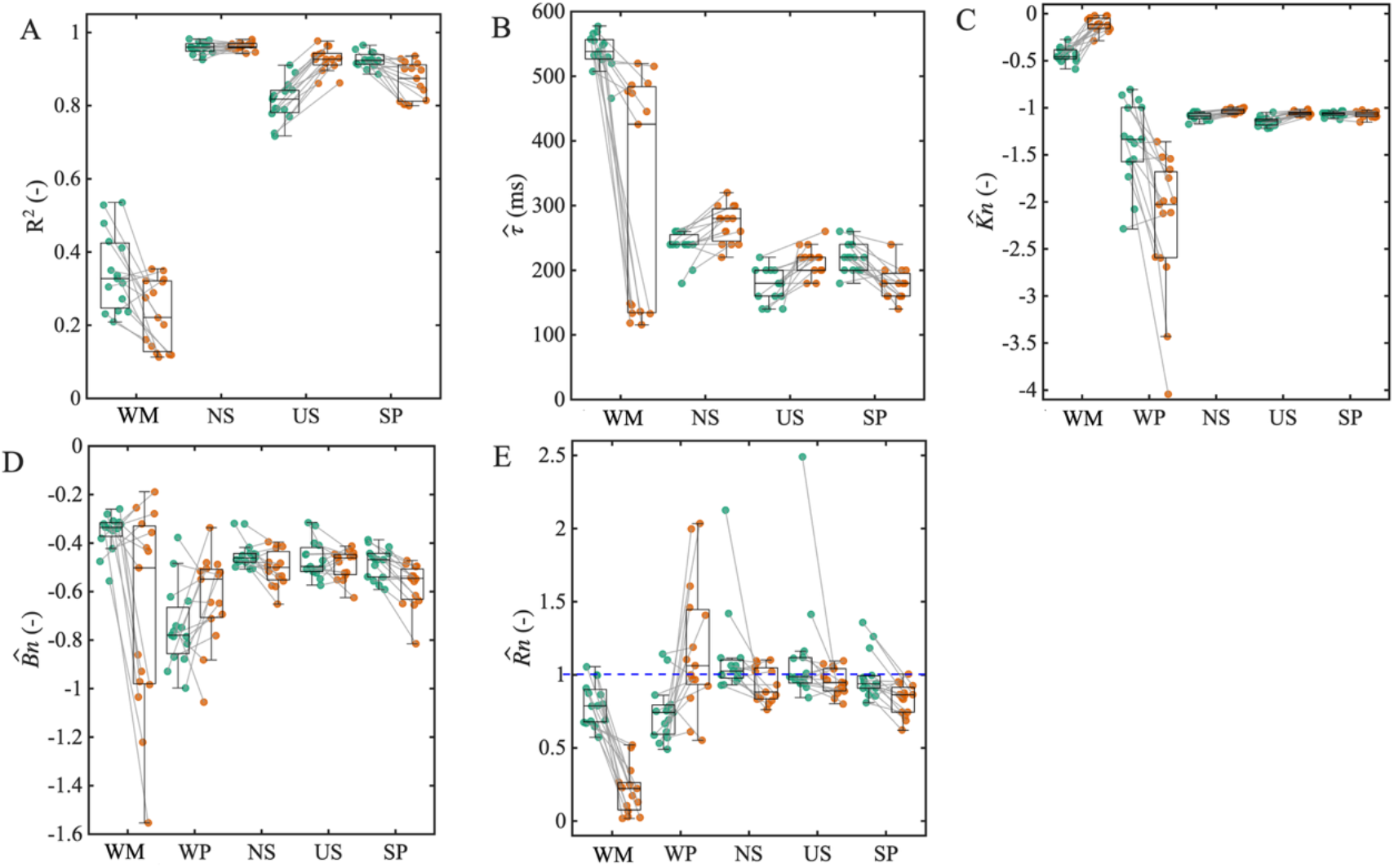
Model fit and estimated stabilizing parameters of all tasks. A) model fit, B) effective delay, C) normalized position gain, D) normalized velocity gain, E) normalized position to velocity gains ratio. Box-plot shows estimated values across all participants, with central line indicating the median, box edges showing the interquartile range, and whiskers representing the non-outlier range. Dots represent individual participant’s data, with green for ML and orange for the AP direction, grey lines connect the data from same participant. The blue dashed line in panel E represent *R*^6^*_n_*=1. X-axis labels denote the following conditions: WM represents the mean values in walking; WP represents the peak values in walking; NS: normal standing; US: unipedal standing; SP: step posture.

**Table 1.**
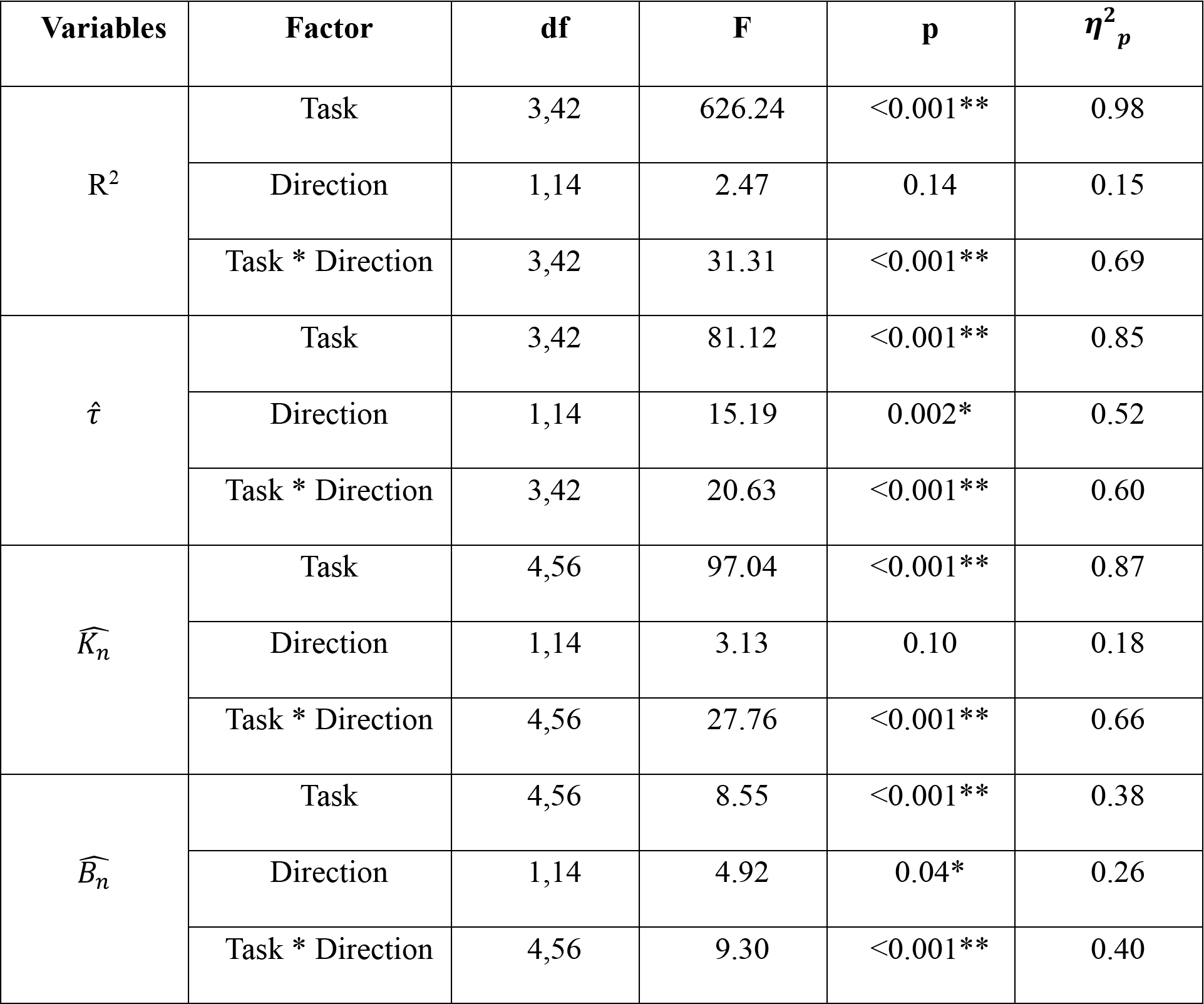
Statistical results of model fit and stabilizing parameters across tasks. For the model fit (*R^2^*), only mean values were used for walking. For the normalized position gains (*K*^6^*_n_*) and velocity gains (*B*^6^*_n_*), both the mean and peak values were used to compare with standing. The asterisks indicating levels of significance, where * indicates p < 0.05 and ** indicates p<0.001.

#### 4.2.1 Model fit

The model fits were low in walking, with mean *R*^2^ values of 0.35 and 0.28 in ML and AP directions, respectively, whereas the mean *R*^2^ values were high in standing, ranging from 0.72 to 0.98 across tasks and directions (Table 1, Fig 3).

Post-hoc simple effects showed that, standing had significantly better model fits than walking in both ML and AP directions (p<0.001). Among the standing tasks, normal standing yielded significantly better model fit than both unipedal standing and step posture (p<0.001). The comparison between unipedal standing and step posture revealed a direction-dependent pattern: step posture had a significantly better model fit in the ML direction (p<0.001), whereas unipedal standing had a significantly better model fit in the AP direction (p=0.02).

There was also direction-dependency within tasks. Walking and step posture showed a significantly better model fit in the ML direction than in the AP direction (p<0.01), whereas unipedal standing showed the opposite pattern, with a better model fit in the AP direction (p<0.001). Normal standing showed no significant difference between directions (p=0.37).

#### 4.2.2 Effective delay

Walking yielded longer effective delays (ML: 538±28ms, AP: 317±180ms) than standing (ranging from 176ms to 239ms across tasks and directions). This difference was significant in the ML direction (p<0.001), while in the AP direction, walking only yielded a longer delay than the step posture (p=0.048). Among standing postures, the pattern differed by direction. In the ML direction, unipedal standing had a significantly shorter delay than both normal standing (p<0.001) and step posture (p<0.001), with no differences between normal standing and step posture (p=0.62). In the AP direction, normal standing had the longest effective delay, followed by unipedal standing, which was significantly longer than the step posture (p=0.017).

For the direction effects, walking and step posture had longer delays in the ML direction than in the AP direction (p<0.001), whereas the opposite pattern was found in normal standing and unipedal standing, with longer delays in the AP direction (p<0.001).

#### 4.2.3 Position gain

Walking yielded a clear phase-dependent modulation of lumped position gains. The mean lumped position gains in walking were lower than the critical stiffness (*K*^^^*_n_*> −1) in both directions and were significantly lower than the position gains in standing (p<0.001). The peak values in walking and values in standing all exceeded their critical values (*K*^^^*_n_*< −1). The peak gains in walking were comparable to standing in the ML direction (p>0.05), but significantly larger than in standing in the AP direction (p<0.001). Among standing tasks, in the ML direction, unipedal standing had significantly larger position gains, followed by normal standing, and both were significantly larger than gains in step posture (p<0.05). In the AP direction, the step posture and unipedal standing had comparable position gains (p=1), both significantly larger than normal standing (p<0.05).

Regarding the direction effects, step posture showed no significant differences between directions (p=0.62), whereas normal standing and unipedal standing had significantly larger position gains in the ML direction (p<0.001). For walking, the mean position gains were significantly larger in the ML direction (p<0.001), while the peak values were significantly larger in the AP direction (p<0.001).

#### 4.2.4 Velocity gain

Similar to the position gains, velocity gains also showed a clear phase-dependency during walking. Velocity gains were below critical damping in all conditions with *_n_*> −1. In the ML direction, the mean velocity gain in walking was significantly lower than the velocity gains in standing (p<0.05), whereas the peak value was significantly larger than in standing (p<0.001). In the AP direction, both mean and peak velocity gains were comparable to those in standing (p>0.05). Among standing postures, no significant differences were observed in the ML direction (p>0.05), whereas in the AP direction, step posture had significantly larger velocity gains than normal standing (p=0.04) and unipedal standing (p=0.02).

No significant directional differences were observed in walking and unipedal standing (p>0.05), whereas the other tasks showed significantly larger velocity gain in the AP direction than in the ML direction (p<0.05).

### 4.3 Ratio of position to velocity gains

The normalized position to velocity gains ratio (*R*^^^*_n_*) was not significantly different from 1 in most conditions (Table 2). Significant deviations from 1 were observed for the mean walking gains in both directions (p<0.001) and the peak walking gains in the ML direction (p<0.001), as well as in normal standing (p=0.04) and step posture in the AP direction (p<0.001).

**Table 2.** Results of one-sample t-tests comparing normalized ratio against 1 across tasks. The asterisks indicating levels of significance, where * indicates p < 0.05 and ** indicates p<0.001. WM: the ratio of mean gains in walking; WP: the ratio of peak gains in walking; NS: normal standing; US: unipedal standing; SP: step posture.

| Tasks | Direction | df | t | p | Cohen's d |
| --- | --- | --- | --- | --- | --- |
| WM | ML | 14 | -5.28 | <0.001** | -1.36 |
|  | AP | 14 | -19.38 | <0.001** | -5.00 |
| WP | ML | 14 | -5.39 | <0.001** | -1.39 |
|  | AP | 14 | 1.59 | 0.14 | 0.41 |
| NS | ML | 14 | 1.47 | 0.16 | 0.38 |
|  | AP | 14 | -2.22 | 0.04* | -0.57 |
| US | ML | 14 | 1.18 | 0.26 | 0.30 |
|  | AP | 14 | -1.80 | 0.09 | -0.46 |
| SP | ML | 14 | -0.32 | 0.76 | -0.08 |
|  | AP | 14 | -5.97 | <0.001** | -1.54 |

We further checked how the ratio changed across the whole stance phase in walking (Fig 4). The ratio in the ML direction remained relatively stable and close to the eigenfrequency in most phases. In the AP direction, the ratio was close to the eigenfrequency only during 55%∼65% of the gait cycle, which coincided with the phases in which the fit of the stabilization model was significant. During the remaining phases the ratio deviated substantially from the eigenfrequency.

**Fig 4.**
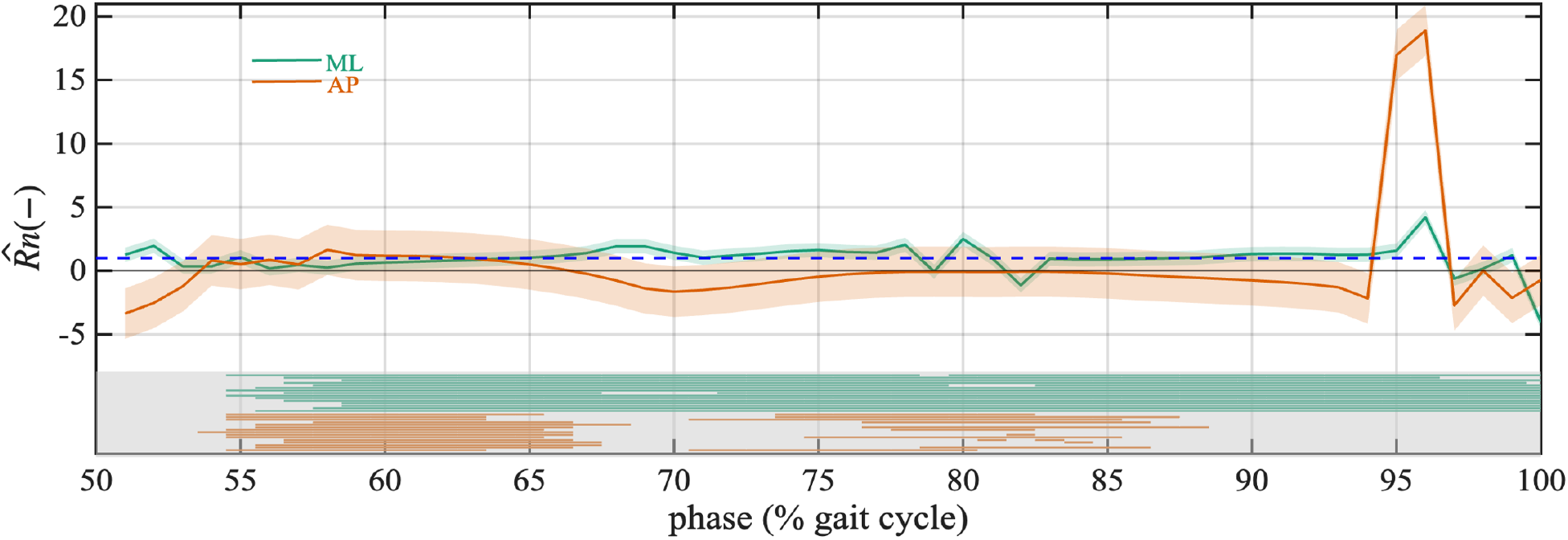
Phase-dependent change in the ratio of position to velocity gains in walking. The green and orange lines represent the averaged results (over participants) in ML and AP direction, respectively. The shaded areas indicate the 95% confidence intervals. The horizontal lines in the grey part of the figure indicate phases in which the stabilization model was significant for each individual participant (p<0.05). Note that the high ratios in the AP direction around 95% correspond with velocity gains close to zero.

## 5. Discussion

In the present study, we fitted a stabilization model to CoM kinematics and ground reaction forces to investigate and compare the lumped stabilizing effects of intrinsic mechanics and feedback control across standing and walking. We found that stabilization of unperturbed standing was well captured by the stabilization model, whereas in unperturbed walking, the stabilization was only partly explained by this model. The lower model fit in walking may reflect, effects of gravity that we could not model using inverted pendulum assumptions, the greater complexity of phase-dependent control, and the additional constraints on force for forward progression besides stabilization.

Despite the fact that the stabilization parameters varied significantly across tasks and directions, all tasks showed under-damped control. Most tasks also had lumped position gains exceeding critical stiffness, except the mean position gains in walking, which were below critical stiffness. The ratio of position to velocity gains remained close to the body’s eigenfrequency predicted by the extrapolated center of mass (XCoM) concept, both in standing and during phases of walking where the stabilization model provided a significant fit.

Our results showed that individual lumped stabilizing parameters are modulated to meet task-dependent demands, whereas the relative control of position and velocity remains consistent whenever stabilization is effective. This consistent weighting between position and velocity contribution reflects a common task-level goal across tasks: maintaining the CoM motion to follow a pendulum-like trajectory.

### 5.1 Utility of the stabilization model in walking

Afschrift et al. described phase-dependent characteristics of feedback responses to perturbations of walking at different speeds, with the responses to perturbations as most pronounced during mid-stance [10]. Our results also showed similar phase-dependent modulation of lumped stabilizing gains, but peak gains were observed during early stance. The observed timing difference can be attributed to the fact that we estimated lumped effects, while the effects of the intrinsic mechanisms are likely to be most pronounced in early stance. Furthermore, methodological differences may also have affected the timing of the peak gains, because we used ground reaction forces as the model output and selected the effective delay based on the variation of gains across the gait cycle, whereas Afschrift et al. [10] used the EMG and ankle torque as model output and used a constant feedback delay.

We found that the effective delay and stabilizing gains differed between ML and AP directions, which may suggest different stability demands and stabilizing strategies between directions. Specifically, the effective delay in the ML direction was longer than in the AP direction. Longer ML delay can be related to larger stability demands [29–32] and more reliance on foot placement [30, 33], while stabilization in the AP direction has been suggested to rely more on the ankle strategy [9, 34], which can take effect during the stance phase and hence at a shorter delay. The overall control of position and velocity in walking, represented by their mean values across the stance phase, showed directional differences: mean position gains were larger in the ML direction, while mean velocity gains were larger in the AP direction. This directional difference was similar to that observed when predicting foot placement based on the CoM state [35], showing a larger contribution of CoM position to ML foot placement and larger contribution of CoM velocity to AP foot placement.

### 5.2 Comparison of stabilization parameters between tasks

#### 5.2.1 Effective delay

As hypothesized, the estimated effective delay was significantly longer in walking than in standing, but only in the ML direction. A longer effective delay in walking was expected, because stabilization is dominated by foot placement and corrections are mainly applied after foot contact [30, 32, 33].

The differences in effective delays between standing postures can be attributed to different biomechanical and neurophysiological characteristics. As noted in the methods section [19], increased intrinsic stiffness leads to a negative bias in effective delays. The observed shorter delays in unipedal standing may thus be explained by an increased intrinsic stiffness due to the higher co-contraction levels [2, 36–40]. In the step posture, its non-neutral ankle angles coincide with higher intrinsic stiffness compared to normal standing, due to the non-linear relationship between ankle angle and elastic parameters [41]. In addition, the geometric configuration of the legs in the step posture with the non-aligned ankle axes will yield ‘structural stiffness’, enhancing overall stiffness. In addition to biasing the estimates of the delay due to differences in intrinsic stiffness, higher muscle activity can be expected to cause a shorter electro-mechanical delay [42], contributing to a shorter effective delay in unipedal standing and step postures.

#### 5.2.2 Position gains

The mean lumped position gains in walking were lower than critical stability (i.e.,*K*^^^*_n_* > −1), whereas the peak values exceeded the critical value. This likely reflects allowing forward progression, while ensuring stability by correcting deviations in CoM position during early stance only. Below critical values during other phases may allow smooth progression while ‘instability’ can be prevented by changes in the base of support (stepping). The lumped position gains in standing exceeded the critical stiffness, which was theoretically expected, as the CoM needs to be tightly controlled over the small base of support to avoid the need to step.

Among standing postures, unipedal standing and step posture had larger lumped position gains than normal standing, which can partly be attributed to their higher intrinsic stiffness as mentioned above. However, feedback control might also contribute to these differences, and it is impossible to separate the intrinsic and feedback contributions without applying external perturbations [8]. Thus, it remains unclear to what extent feedback mechanisms contribute to the differences in lumped gains. Interestingly, lumped position gains in unipedal standing were approximately twice the per leg values observed in normal standing, despite the smaller base of support and the reduction in number of muscles that can contribute to the corrective forces. This indicates that the control system compensates for the reduced mechanical stability by increasing the lumped gains per leg, reflecting a flexible adjustment of the combined stabilizing mechanisms. Similarly, it is reasonable to speculate that the feedback control in the step posture might be decreased due to the additional structural stiffness.

#### 5.2.3 Velocity gains

The lumped velocity gains revealed an under-damped control in both walking and standing. Previous studies have also described human upright standing as underdamped [43, 44]. While it does allow some oscillatory behavior around the target state, underdamped control causes faster return to states close to the target state than critically damped control.

#### 5.2.4 Ratio of position to velocity gains

According to the XCoM concept, pendular movement requires the ratio of position to velocity gains to equal the system’s eigenfrequency, yielding a normalized ratio (*R*^^^*_n_*) of 1. This is mainly supported by standing results, with the *R*^^^*_n_* did not differ significantly form1 in most tasks, and although the differences were significant in normal standing and step posture in the AP direction, the values remained numerically close to 1. In walking, although the ratio of mean gains in both directions and of peak gains in the ML direction differed significantly from 1, *R*^^^*_n_* remained close to 1 whenever the stabilization model was significant, suggesting that pendular-like movement control is preserved when the stabilization control is pronounced.

## 6. Limitations

Several limitations should be noted. First, we modeled the human body as a single inverted-pendulum in standing to calculate the corrected forces. However, this does not fully capture the multi-segmented nature of the human body, which may have introduced errors in force decomposition. However, the fact that the ratio of position to velocity gains in most standing condition was close to eigenfrequency suggests that this simplification does not substantially affect the estimation of overall stabilizing mechanisms in the current study. Second, we used the total ground reaction forces, which include some effects of gravity, rather the corrected forces to estimate the stabilizing parameters in walking. Also, we applied different methods to select the effective delay and corresponding lumped gains between standing and walking, which may lead to some bias when comparing the stabilizing parameters. In addition, walking may introduce more soft tissue artifacts in the measurement of kinematics, potentially affecting the parameter estimation. Nonetheless, the estimated parameters aligned well with the task-specific characteristics. Furthermore, while the lumped model captures the overall stabilizing effects, separating intrinsic and feedback contributions would provide more detailed information, but this requires perturbation-based studies [8]. Finally, we examined only one gait speed in this study, whether and how the observed feedback strategy will change with speed needs further examination.

## 7. Conclusion

Our results showed that lumped stabilizing parameters: effective delay, lumped position and velocity gains, differed significantly across standing and walking tasks and mediolateral and anteroposterior directions. While the individual parameters differed significantly, the ratio of position to velocity gains remained close to the body’s eigenfrequency whenever stabilization was pronounced, reflecting a common task-level goal: keeping the CoM on a nominal pendulum-like trajectory.

## Acknowledgements

The authors thank all the participants for their participation in this study.

## Supporting information

**S1 Appendix.**

**S1 Table.** Estimated stabilization results in walking and standing (mean±SD).

**S1 Figure.** Normalized time series data during walking in ML (panel A) and AP direction (panel B) from one typical participant.

## Notes

https://zenodo.org/records/22304652

### Competing Interest Statement

The authors have declared no competing interest.

### Summary of Updates

There was an error in the code for processing the data on walking, code has been updated in Zenodo. All affected results, Fig 2-4, Table 1 and 2, and corresponding text have been revised.Supplemental files have been updated.

### Funding

- China Scholarship Council

