## Supplementary material for "Different stabilizing mechanisms but a common task-level aim in standing and walking": S1 Appendix

1. Reliability of estimated lumped parameters

After obtaining the effective delay ($\hat{\tau}$), lumped position ($\hat{K}$) and velocity gains ($\hat{B}$) for all 5 trials and all tasks (**Fig 1**), we evaluated the relative and absolute reliability of these parameters by calculating the intra-class correlation coefficients (ICC) and standard error of measurement (SEM) [1-3]. Data from one participant (S04) were excluded for the ICC and SEM calculations for normal standing, because this participant had only 4 trials. Because we used the mean values of repeated trials as the estimated values for each participant, the ICC was calculated using one-way random-effect model as defined by equation (1):

$$\begin{aligned} ICC\left( 1,k \right)=\frac{{MS}_{B}-{MS}_{W}}{{MS}_{B}}\#\left( 1 \right) \end{aligned}$$

where the ${MS}_{B}$ is the between subject mean square, ${MS}_{W}$ is the within subject mean square, $k$ is the number of trials which was 5 in this study.

The SEM was calculated using equation (2) $\begin{aligned} SEM=\sqrt{{MS}_{w}/k}\#\left( 2 \right) \end{aligned}$

The ICC and SEM values of estimated stabilization parameters are shown in **Table 1**. ICC values less than 0.5 were regarded as indicating poor reliability, values between 0.5 and 0.75 were regarded as indicating moderate reliability, values between 0.75 and 0.9 were regarded as indicating good reliability, and values higher than 0.9 were regarded as indicating excellent reliability[3].

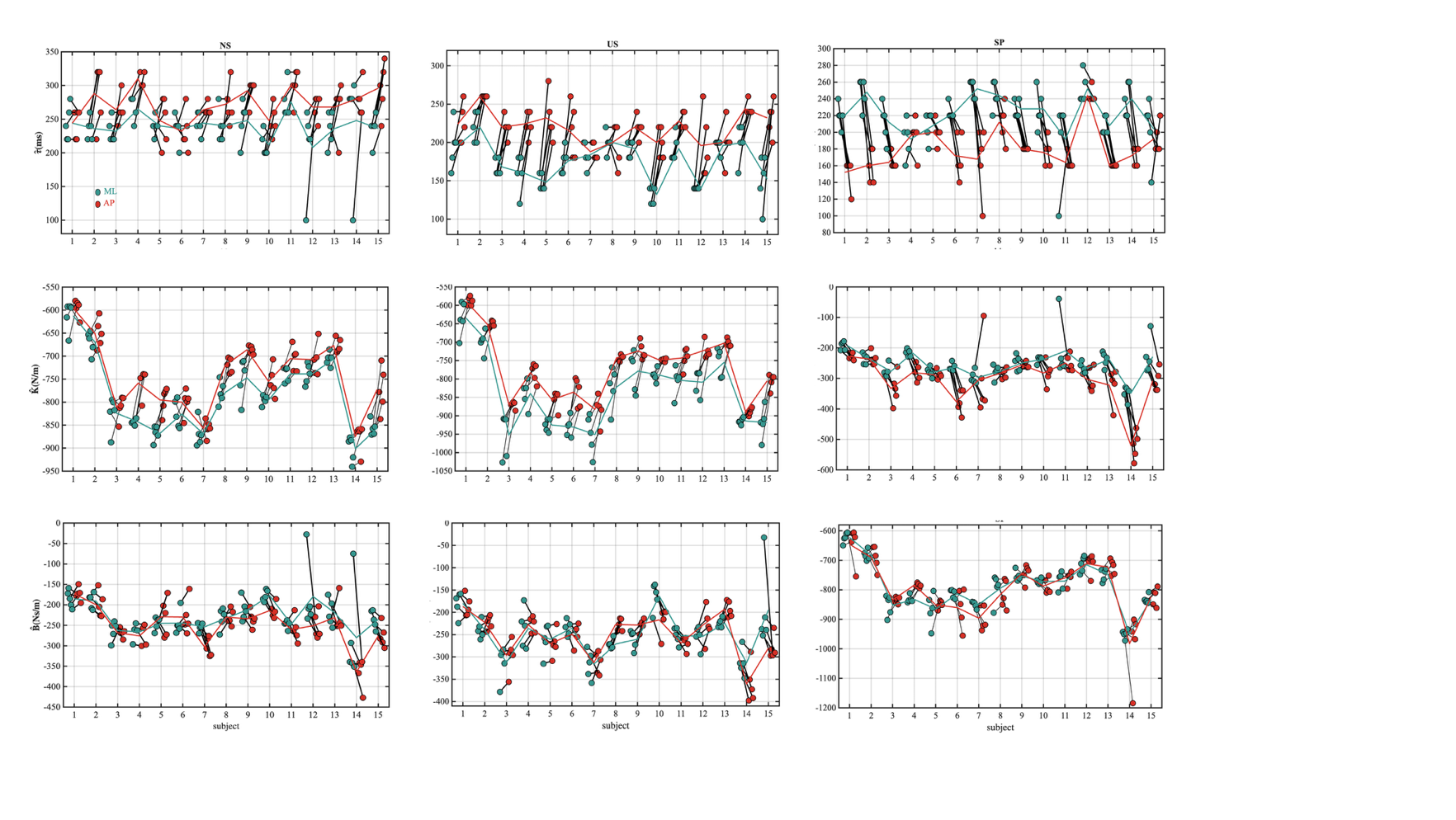

**Fig 1. Estimated stabilization parameters across all trials, tasks, and participants.** The dots represent values of separate one-minute trials, the lines represent the mean values of all trials for all participants. NS: normal standing; US: unipedal standing; SP: step posture.

**Table 1. ICC and SEM results for estimated stabilization parameters.**

| Parameter | Direction | Task | ICC (1,5) | SEM |
| --- | --- | --- | --- | --- |
| $\hat{\tau}$ | ML | NS | -0.07 | 15 |
|  |  | US | 0.90 | 8 |
|  |  | SP | 0.75 | 10 |
|  | AP | NS | 0.62 | 13 |
|  |  | US | 0.71 | 11 |
|  |  | SP | 0.88 | 8 |
| $\hat{K}$ | ML | NS | 0.98 | 12 |
|  |  | US | 0.97 | 18 |
|  |  | SP | 0.98 | 12 |
|  | AP | NS | 0.98 | 12 |
|  |  | US | 0.99 | 10 |
|  |  | SP | 0.94 | 21 |
| $\hat{B}$ | ML | NS | 0.65 | 20 |
|  |  | US | 0.88 | 16 |
|  |  | SP | 0.85 | 15 |
|  | AP | NS | 0.92 | 13 |
|  |  | US | 0.94 | 12 |
|  |  | SP | 0.92 | 20 |

NS: normal standing; US: unipedal standing; SP: step posture.

The figure of all estimated parameters (**Fig 1**) shows that the estimated delays had less variance between participants than within participants, and the mean values of all trials were relatively similar between participants. The lumped position gains were highly consistent within participants and differed significantly between participants. In contrast, the lumped velocity gains showed somewhat greater within-participant variance but smaller between-participant variance than the lumped position gains.

Consistent with the patterns observed in **Fig 1**, the ICC and SEM results showed that the lumped position gains had the highest reliability, followed by the lumped velocity gains, whereas the effective delays showed lower reliability than the gain estimates. Specifically, the effective delays ($\hat{\tau}$) showed good reliability in unipedal standing and step posture, but poor and moderate reliability in normal standing in the ML and AP directions, respectively. However, the SEM values across all conditions were smaller than 20ms, which was the minimal interval used for delay estimation. This indicates that the lower ICC of the effective delay is a consequence of the low variability between participants. The lumped position gains ($\hat{K}$) showed excellent reliability across all conditions, and the corresponding SEM values were relatively low. The lumped velocity gains ($\hat{B}$) across all conditions showed good to excellent reliability, with the exception of normal standing in the ML direction, which showed moderate relative reliability but low SEM values.

These results indicated that the estimated lumped gains were reliable and stable across trials. The lower ICCs of the effective delay and in some cases for the velocity gains may reflect a mathematical trade-off between delays and lumped gains. In the following section, we further report on tests of the model’s identifiability.

2. Identifiability of stabilization parameters by the proposed model

We first examined whether a trade-off between delays and lumped gains exists by checking how the gains changed with imposed effective delays (**Fig *2***). Results (**Fig *2***) shows that there is indeed a trade-off between parameters: shorter effective delays are accompanied by larger position gains and smaller velocity gains.

Then we assessed the identifiability of the stabilization parameters under such a trade-off: we calculated the gains ($\hat{K}$and $\hat{B}$) for specific delays ($\hat{\tau}$): ranging from 20ms to 400ms with an interval of 20ms and calculated the ICC and SEM of the lumped gains ($\hat{K}$and $\hat{B}$) for each selected delay. The model fit and estimated $\hat{K}$and $\hat{B}$ for all values of $\hat{\tau}$ are shown in **Fig 3**. To better understand how the trade-off affects the estimated values, we plotted the variability of estimated $\hat{K}$and $\hat{B}$as function of effective delay $\hat{\tau}$, as shown in **Fig 4**.

**Fig 3** shows that the lumped gains substantially varied with delays. **Fig 4** further shows that while the position gains decreased monotonically, the variation was small, less than 25% of the mean value. Velocity gains decreased strongly with increasing delays until the delay reached 80ms, after which there was small variation relative to the mean value.

Consistent with these results, **Fig 5** show that the estimated velocity gains are reliable (ICC>0.75) when the delay is larger than 80ms, while the SEM values show consistent estimates for delays over 140~160ms.

In conclusion, these results show that there is indeed trade-off between parameters, while this effect is quite small for delays larger than 80ms (which was what we found). Thus, using our approach, we can reliably obtain the effective delay and lumped stabilization gain.

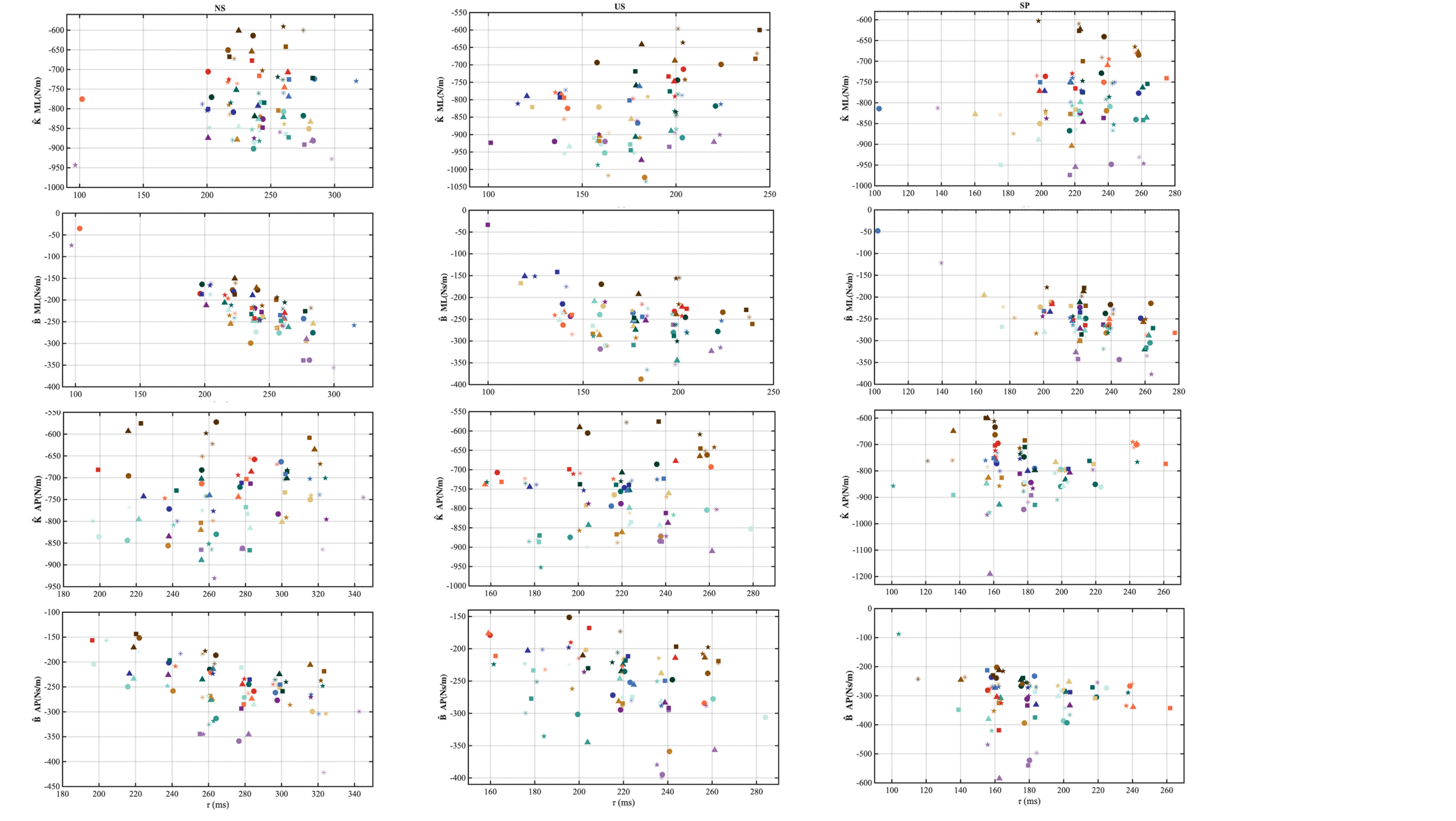

**Fig 2. Stabilization gains plotted against estimated effective delays across all trials, tasks, and participants.** Different colors represent individual participants. Each dot represents the value from a separate one-minute trial with different marker shapes indicate different trials. NS: normal standing; US: unipedal standing; SP: step posture.

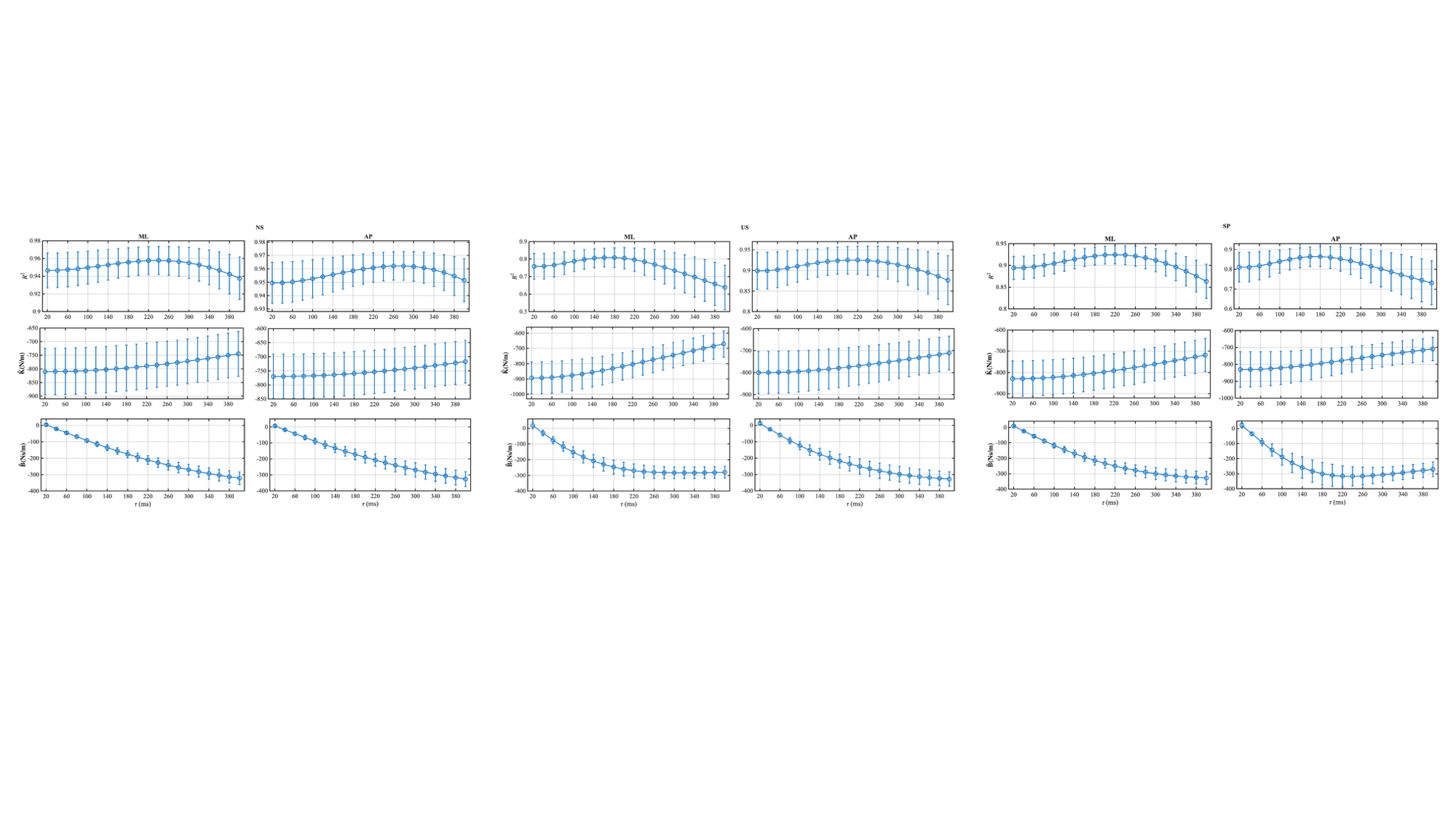

**Fig 3. Model fits and estimated stabilization gains under selected delays.** NS: normal standing; US: unipedal standing; SP: step posture.

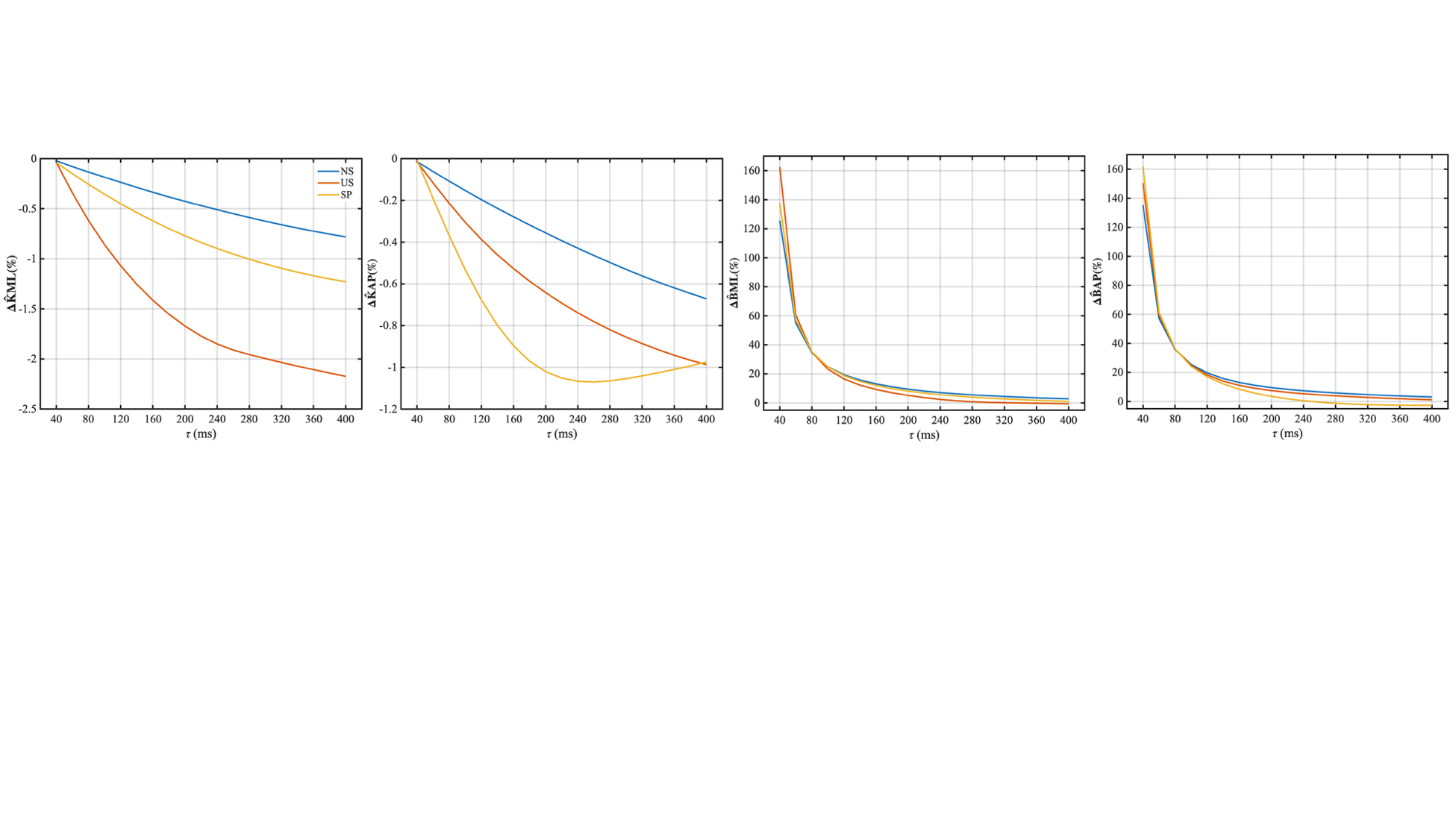

**Fig 4. Variability of** $\hat{\mathbf{K}}$**and** $\hat{\mathbf{B}}$**as function of** $\hat{\boldsymbol{\tau}}$ **for different standing postures.** The x-axis represents the specific delay τ (ms). The y-axis shows the relative change expressed as a percentage of the parameter value at the corresponding delay. NS: normal standing; US: unipedal standing; SP: step posture.

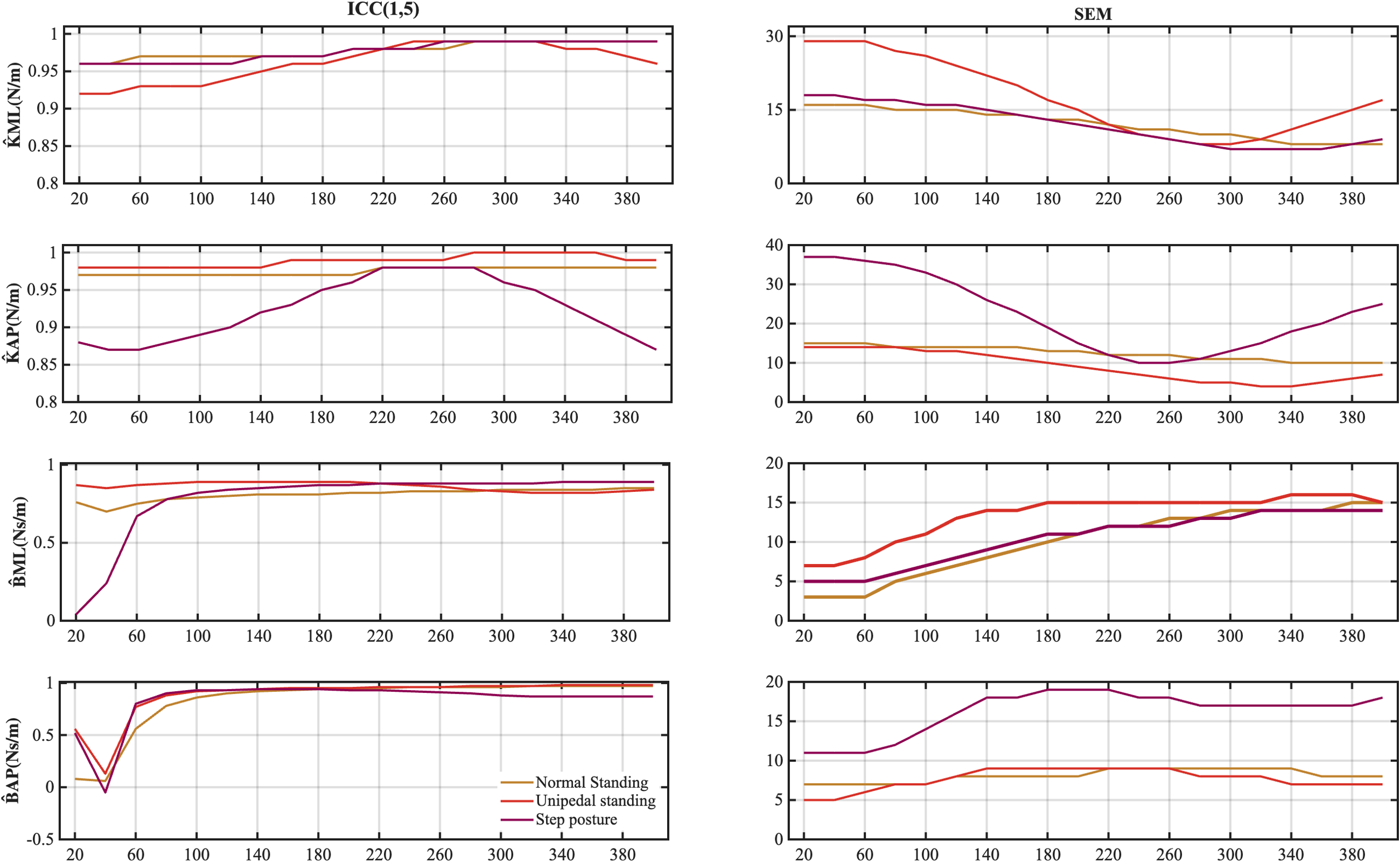

**Fig 5. ICC and SEM values under selected delay.** NS: normal standing; US: unipedal standing; SP: step posture.
