## Supplementary material for "Different stabilizing mechanisms but a common task-level aim in standing and walking": S1 Figure

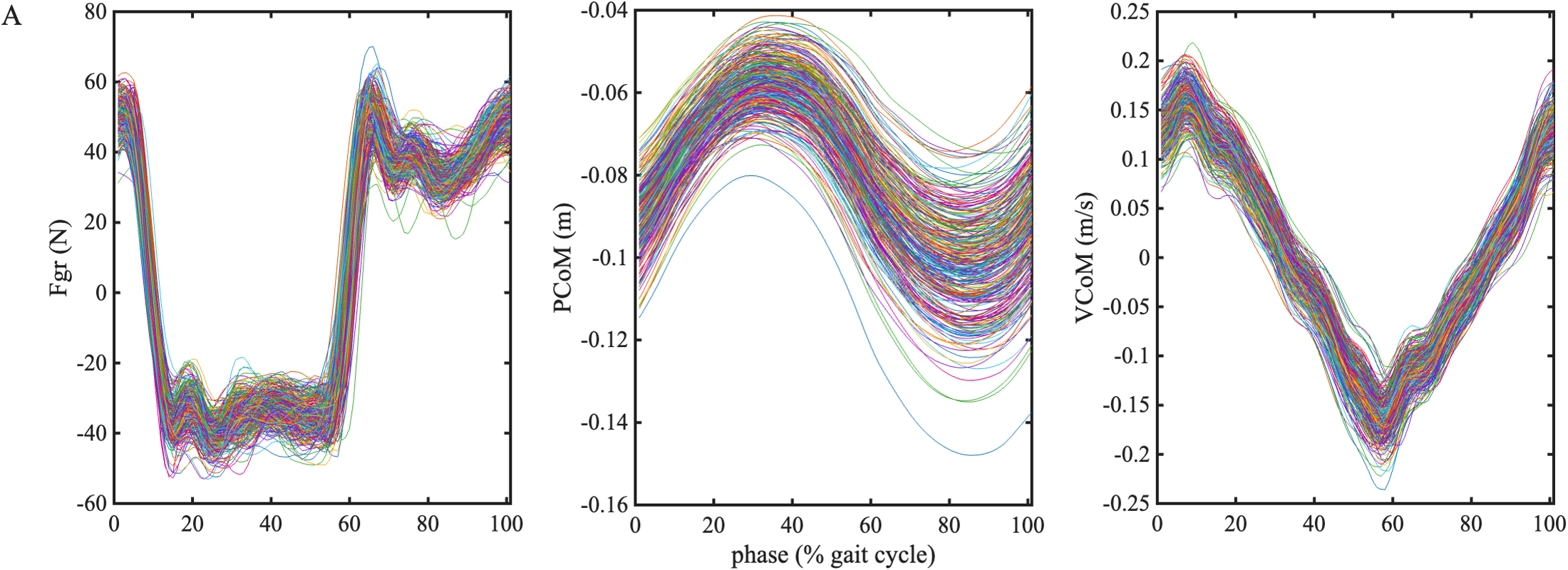

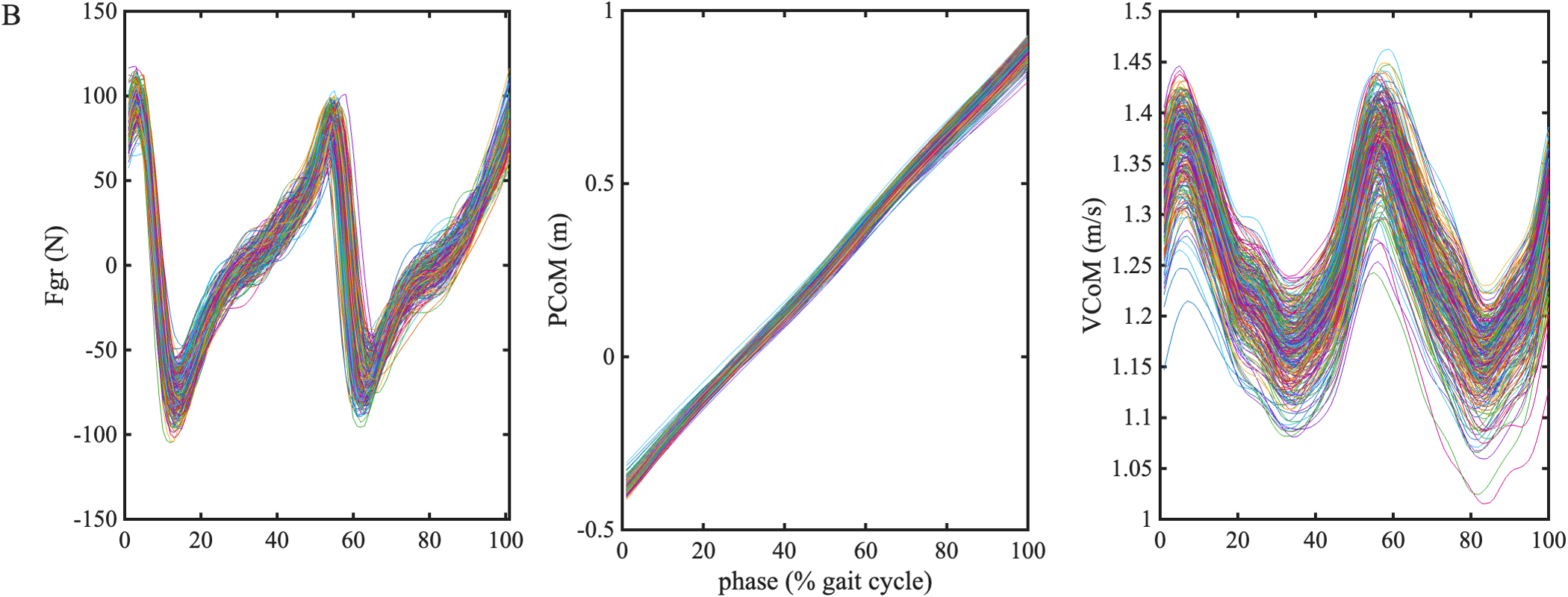

**S1 Fig. Normalized time series data during walking in ML (panel A) and AP direction (panel B) from one typical participant.** From left to right: ground reaction forces (Fgr), CoM position (PCoM) and velocity (VCoM) across the gait cycle (normalized based on right heel strike). Different colors represent different strides. Note the PCoM data were calculated from the raw PCoM data after re-referencing them to the mean trailing foot position during the 15%~45% phase of each stride.
