## Supplementary material for "Different stabilizing mechanisms but a common task-level aim in standing and walking": S1 Table

**S1 Table. Estimated stabilization results in walking and standing (mean±SD).**

| **Variables** | **Directions** | **WM** | **WP** | **NS** | **US** | **SP** |
| --- | --- | --- | --- | --- | --- | --- |
| R^2^ (-) | ML | 0.35±0.11 |  | 0.96±0.02 | 0.81±0.05 | 0.92±0.02 |
|  | AP | 0.23±0.09 |  | 0.96±0.01 | 0.93±0.03 | 0.87±0.05 |
| $\hat{\tau}$ (ms) | ML | 538±28 |  | 239±22 | 176±25 | 221±23 |
|  | AP | 317±180 |  | 271±28 | 216±22 | 179±24 |
| $\hat{K_{n}}$ (-) | ML | -0.44±0.08 | -1.38±0.44 | -1.09±0.04 | -1.15±0.05 | -1.07±0.03 |
|  | AP | -0.11±0.07 | -2.23±0.74 | -1.03±0.02 | -1.06±0.02 | -1.07±0.03 |
| $\hat{B_{n}}$ (-) | ML | -0.36±0.08 | -0.75±0.16 | -0.45±0.06 | -0.47±0.08 | -0.48±0.06 |
|  | AP | -0.69±0.42 | -0.62±0.18 | -0.50±0.07 | -0.49±0.06 | -0.57±0.09 |
| $\hat{R_{KB}}$ (-) | ML | 0.80±0.15 | 0.74±0.19 | 1.12±0.30 | 1.12±0.40 | 0.99±0.16 |
|  | AP | 0.21±0.16 | 1.18±0.44 | 0.93±0.12 | 0.96±0.09 | 0.83±0.11 |

WM represents the mean values in walking; WP represents the peak values in walking; NS: normal standing; US: unipedal standing; SP: step posture.
